# Exploiting HSD17B11-dependent dialkynylcarbinols cytotoxicity for facile CRISPR/Cas9-based gene inactivation

**DOI:** 10.64898/2026.05.13.724824

**Authors:** Bastien Dumais, Madeleine Bossaert, Patrick Seigneur, Alexandrine Rozié, Sophie Gasmi, Luce Fontanié--Roger, Maëlle Caroff, Valérie Maraval, Vania Bernardes-Génisson, Dennis Gomez, Philippe Frit, Stéphanie Ballereau, Yves Génisson, Sébastien Britton

## Abstract

Several approaches have been developed to improve the efficiency of CRISPR/Cas9-based genome editing, including the co-inactivation of a gene whose loss confers resistance to a cytotoxic compound, thereby enabling enrichment of successfully edited cells. Here, we show across multiple cell lines that inactivation of HSD17B11, a non-essential member of the Short-chain Dehydrogenase/Reductase (SDR) superfamily, confers strong resistance (29-to 131-fold) to a Phenyl diAlkynylCarbinol compound (PAC) in both human and mouse cells, without affecting cell viability or proliferation. We demonstrate that co-inactivation of HSD17B11 followed by PAC selection can be used to rapidly identify efficient guide RNAs targeting a gene of interest and to readily isolate clones inactivated for one or multiple genes. Altogether, these results establish a simple and efficient experimental strategy for generating knockout cells by using PAC selection to enrich for successfully edited cells.

## Introduction

The simplest approach for gene inactivation using CRISPR/Cas9 relies on unfaithful DNA repair of a DNA double-strand break induced by Cas9 at the target locus resulting in insertions or deletions at the cleavage site that can disrupt gene function. More advanced strategies have been developed to improve editing specificity or control the resulting genetic alteration, for example by using paired proximal nicks generated by the *Streptococcus pyogenes* Cas9-D10A mutant (referred to as Cas9 nickase), or by directly introducing stop codons using base-editors or prime-editors Cas9 fusion proteins **(Billon et al., 2017; Nambiar et al., 2022; Pacesa et al., 2024)**. Despite these approaches, a major bottleneck remains the low frequency of the event(s) of interest within the cell population.

Multiple strategies have been developed to circumvent this limitation, which include i) improved delivery, for example with viral particles **(Djamshidi et al., 2026; Lyu et al., 2019)**, ii) stable expression of Cas9 in target cells, or iii) enrichment of successfully edited cells. For example, co-expressing Cas9 with a fluorescent protein enables Fluorescence-Activated Cell Sorting (FACS) to enrich for transfected cells in which Cas9 is successfully expressed, but requires the proper facilities **(Ran et al., 2013)**. An alternative strategy readily applicable consists of co-editing a gene whose alteration confers resistance to a cytotoxic compound. Subsequent drug selection enriches for cells in which Cas9 activity was sufficient to edit the selection gene and, consequently, increases the probability of recovering cells co-edited at the gene of interest (GOI). When the guide RNA (gRNA) targeting the GOI is provided in excess relative to the gRNA targeting the selection gene, a large proportion of drug-resistant cells can also carry the desired alteration at the GOI, provided that the corresponding gRNA is efficient.

A widely used compound for this type of enrichment is ouabain **(Agudelo et al., 2017)**. Ouabain, a cardiotonic steroid, is a potent inhibitor of the human Na^+^/K^+^-ATPase, a ubiquitous and essential ion transporter responsible for maintenance of the electrochemical gradients of Na^+^ and K^+^ across the plasma membranes of animal cells. Ouabain is cytotoxic towards human cells and introduction of specific mutations in the ATP1A1 gene coding for the Na^+^/K^+^ ATPase makes cells resistant to this drug **(Croyle et al., 1997)**. For example, gRNAs targeting ATP1A1 exon 4 can be used to generate in frame deletions within the first extracellular loop of ATP1A1 (Q118-N129) conferring a ∼20-fold resistance exploitable to enrich for co-edited genes **(Agudelo et al., 2017)**. Combinations of point mutations (Q118R+N129D or Q118D+N129R) can also be introduced using single-stranded oligodeoxynucleotides (ssODN) donors to confer >2000-fold resistance to ouabain **(Agudelo et al., 2017)**. This high level of resistance was exploited for sequential editing of multiple genes by first introducing a mutation (T804N, exon 17) conferring a ∼20-fold resistance to enable editing a first gene, followed by a second mutation (Q118R, exon 4) conferring maximal resistance for editing of a second gene **(Levesque et al., 2022)**. This makes the selection of ATP1A1 edited cells an attractive strategy. However, it also has some drawbacks, calling for complementary approaches. First, ATP1A1, the α1 subunit of the Na^+^/K^+^-ATPase, is a widely expressed essential plasma-membrane pump that maintains ionic, osmotic and electrical homeostasis. Because it establishes the transmembrane Na^+^ gradient, even modest changes in pump activity can alter secondary transport processes, including Na^+^-coupled amino-acid and glucose uptake, intracellular pH regulation through Na^+^/H^+^ exchange, and Na^+^/Ca^2+^ exchange, thereby affecting Ca^2+^-dependent cellular processes, including response to multiple drugs **(Contreras et al., 2024; Gagnon and Delpire, 2020)**. Accordingly, ATP1A1 mutations are responsible for several human genetic diseases such as primary aldosteronism and Charcot-Marie-Tooth disease **(Biondo et al., 2021)**. These pleiotropic functions raise the question of the impact of the mutations conferring resistance to ouabain on cell physiology, especially when relying on loop deletions of variable sizes. Finally, multiple species, including rodents, are not sensitive to ouabain and related compounds, making this approach unsuitable in these organisms **(Lingrel, 2010)**.

We recently discovered that the high enantiospecific cytotoxicity of specific chiral alkynylcarbinol lipids, such as DiAlkynylCarbinols (DACs) and their improved congeners Phenyl diAlkynylCarbinols (PACs), is strictly dependent on the catalytic activity of the intracellular HSD17B11 enzyme (**(Bossuat et al., 2023; Demange et al., 2022), Fig. 1A**). HSD17B11 (also known as 17β-hydroxysteroid dehydrogenase type-11, SDR16C2, PAN1B, DHRS8, or retSDR2) is a non-essential protein, ubiquitously expressed in humans and member of the Short-chain Dehydrogenase/Reductase (SDR) superfamily. It localizes to the endoplasmic reticulum (ER) and lipid droplets where it uses NAD^+^ to catalyze oxidation of the C17 carbinol center of androstan-3-alpha,17-beta-diol to generate androsterone, a weak androgen **(Brereton et al., 2001; Horiguchi et al., 2008)**. Mice knock-out for HSD17B11 are viable and show minor phenotypes evocative of increased androgen levels (e.g. increased heart weight, **(Mouse Genome Informatics, 2024)**). We previously established that DAC and PAC are enantiospecifically converted within cells into protein-reactive DiAlkynylCarbinones (DACones) and Phenyl diAlkynylCarbinones (PACones), respectively, by the catalytic activity of HSD17B11. DACones and PACones react with multiple essential intracellular proteins resulting in an acute proteotoxic stress leading to rapid apoptotic cell death (**Fig. 1A**). As compared to DACs, PACs represent an improved generation that shows enhanced stability in biological media and allows easier pharmacomodulation than the original DACs **(Bossuat et al., 2023)**. While (*S*)-PACs are highly cytotoxic, (*R*)-PACs are basically innocuous at the same concentration **(Bossuat et al., 2023)**, allowing the use of the racemic mixture, easier to prepare as compared to enantioenriched compounds, to achieve HSD17B11-dependent cytotoxicity.

**Figure 1.**
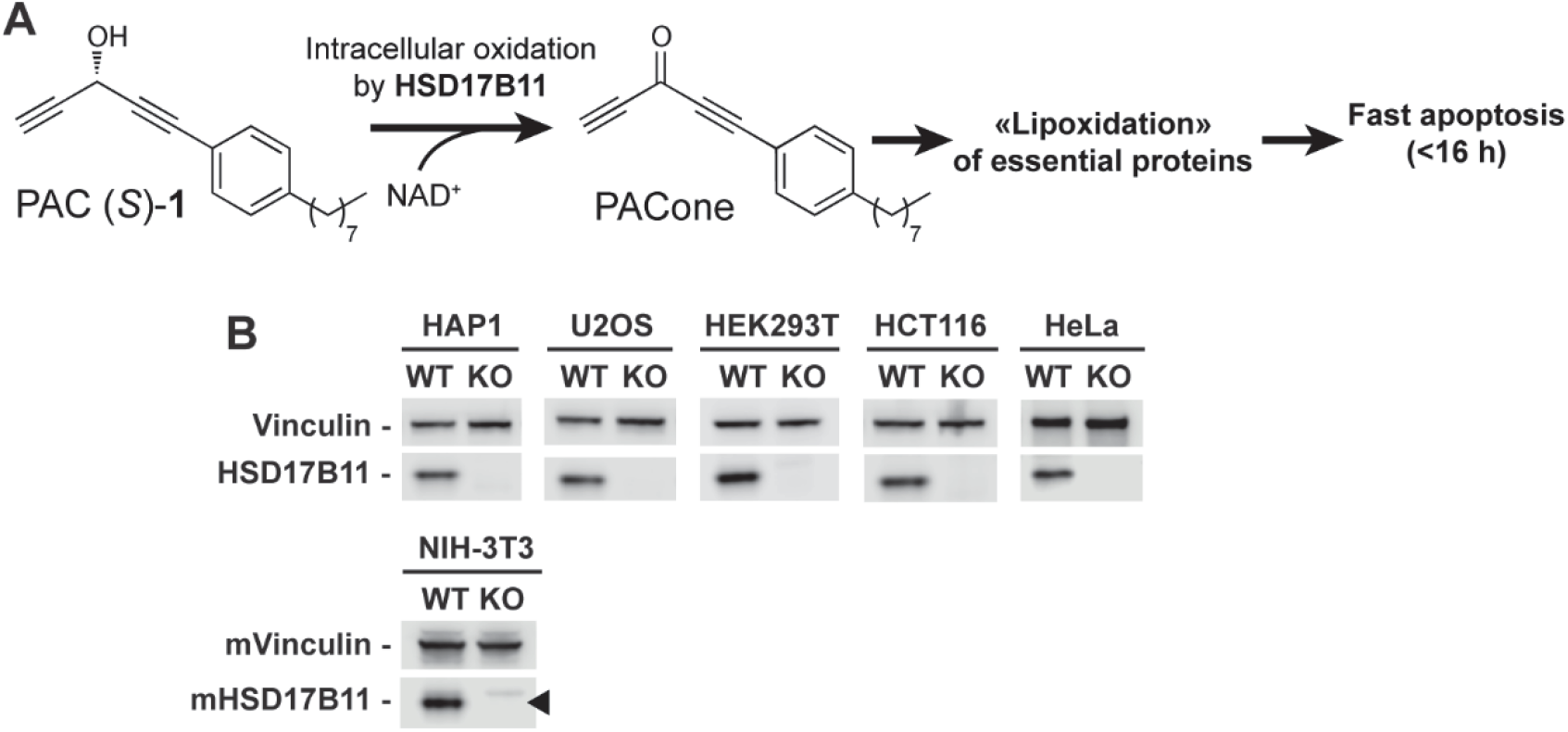
(**A**) Scheme depicting the enantiospecific bioactivation of (S)-PAC **1** by the intracellular SDR enzyme HSD17B11 into the cytotoxic PACone. PACones are highly reactive electrophiles that, when produced in cells, covalently modify multiple proteins including essential factors involved in protein quality control. This leads to an acute ER stress and fast induction of apoptosis. (**B**) Analysis by immunoblotting of HSD17B11 levels in each cell line, including WT and an isolated clone. Vinculin was used as a loading control. The black arrows indicate the band corresponding to HSD17B11.

Here, we exploit the strong resistance to PACs conferred by HSD17B11 inactivation as a selection strategy for CRISPR/Cas9-mediated gene knockout in mammalian cells. We show in multiple human cell lines and in mouse NIH-3T3 how inactivation of HSD17B11 by CRISPR/Cas9 confers a strong resistance to a racemic PAC. Then we illustrate, by targeting TOP2B in human and mouse cells, how co-edition of HSD17B11 and selection with a racemic PAC can be used to identify efficient gRNAs and isolate clones inactivated for the GOI. We also show that this approach allows several GOIs to be inactivated simultaneously. These results establish HSD17B11/PAC selection as a simple approach to facilitate CRISPR/Cas9-based gene inactivation in both human and mouse cells.

## Results

### Inactivation of HSD17B11 using CRISPR/Cas9 confers cellular resistance to a racemic PAC

For this study, we selected the racemic form of PAC (*S*)-**1** depicted in **Fig. 1A (Bossuat et al., 2023)**, which represents the prototype of this family. We transfected various representative laboratory cell lines with the pCAG-eSpCas9-2A-GFP plasmid coding for the *S. pyogenes* Cas9 K848A K1003A R1060A variant, which displays reduced off-target editing **(Slaymaker et al., 2016)** and which co-expresses a gRNA against human or mouse HSD17B11. Six days after transfection, 250 000 cells were seeded in large dishes and selected the day after with a toxic concentration of PAC (2 and 8 µM for human and mouse cells, respectively). Individual clones were isolated and analyzed by immunoblotting using a specific antibody against HSD17B11 to confirm loss of HSD17B11 expression. One clone for each cell line was selected and used for further studies (**Fig. 1B**).

For each cell line, the IC_50_ of the racemic PAC **1** was established for the original cell line and its HSD17B11-KO derivative using sulforhodamine B assays (SRB) after a 72 h treatment with serial 2-fold dilutions (**Table 1**, see **Supplementary Fig. 1** for viability and proliferation curves).

**Table 1.** Summary of doubling time, PAC cytotoxicity, and selection concentration for each cell line.

| Cell line | HAP1 |  | U2OS |  | HEK293T |  | HCT116 |  | HeLa |  | NIH-3T3 |  |
| --- | --- | --- | --- | --- | --- | --- | --- | --- | --- | --- | --- | --- |
|  | WT | KO B11 | WT | KO B11 | WT | KO B11 | WT | KO B11 | WT | KO B11 | WT | KO B11 |
| Doubling Time (h) | 22.32 | 21.5 | 25.03 | 24.58 | 19.55 | 18.43 | 22.65 | 23.47 | 15.69 | 17.73 | 29.17 | 31.13 |
| IC<sub>50</sub> (μM) | 0.1 | 6.28 | 0.12 | 13.44 | 0.14 | 4.02 | 0.13 | 17 | 0.36 | 13.56 | 1.25 | > 50 |
| Resistance Factor | 63 |  | 112 |  | 29 |  | 131 |  | 38 |  | > 40 |  |
| Concentration used for selection (μM) | 2 |  | 2 |  | 2 |  | 2 |  | 2 |  | 8 |  |

HSD17B11 KO cell lines showed 29-to 131-fold higher resistance to PAC **1** as compared to corresponding WT cell lines. In addition, we monitored proliferation of the parental cell lines and their HSD17B11-KO derivatives over 3 days and found no significant effect of HSD17B11 inactivation on the doubling time of any of the cell lines analyzed. These data indicate that HSD17B11 is dispensable for cell growth and viability both in human and mouse cells. This conclusion is also supported by the Depmap data **(Arafeh et al., 2025)**, in which HSD17B11 stands out as a non-essential gene in all the 1,186 cell lines analyzed.

### Co-inactivation of HSD17B11 along with selection with PAC can be exploited to identify efficient gRNAs against a GOI

We reasoned that selection with the racemic PAC could be used to enrich a cell population transfected with plasmid expressing CRISPR/Cas9 targeting a GOI if a smaller amount of plasmids expressing CRISPR/Cas9 complexes against HSD17B11 were included. Co-editing of HSD17B11 would then allow the use of the racemic PAC (**Fig. 1A**) to enrich for cells in which CRISPR/Cas9 editing was efficient following transfection (**Fig. 2A**). Using higher amounts of CRISPR/Cas9 complexes targeting the GOI relative to the selection gene increases the probability that resistant cells will be co-edited at the GOI. As a proof of concept, we focused on the TOP2B gene, coding for the DNA topoisomerase 2 beta protein, which is dispensable for cell viability of human and mouse cell lines **(Arafeh et al., 2025; DepMap, 2026c; Uuskula-Reimand and Wilson, 2022)**.

**Figure 2:**
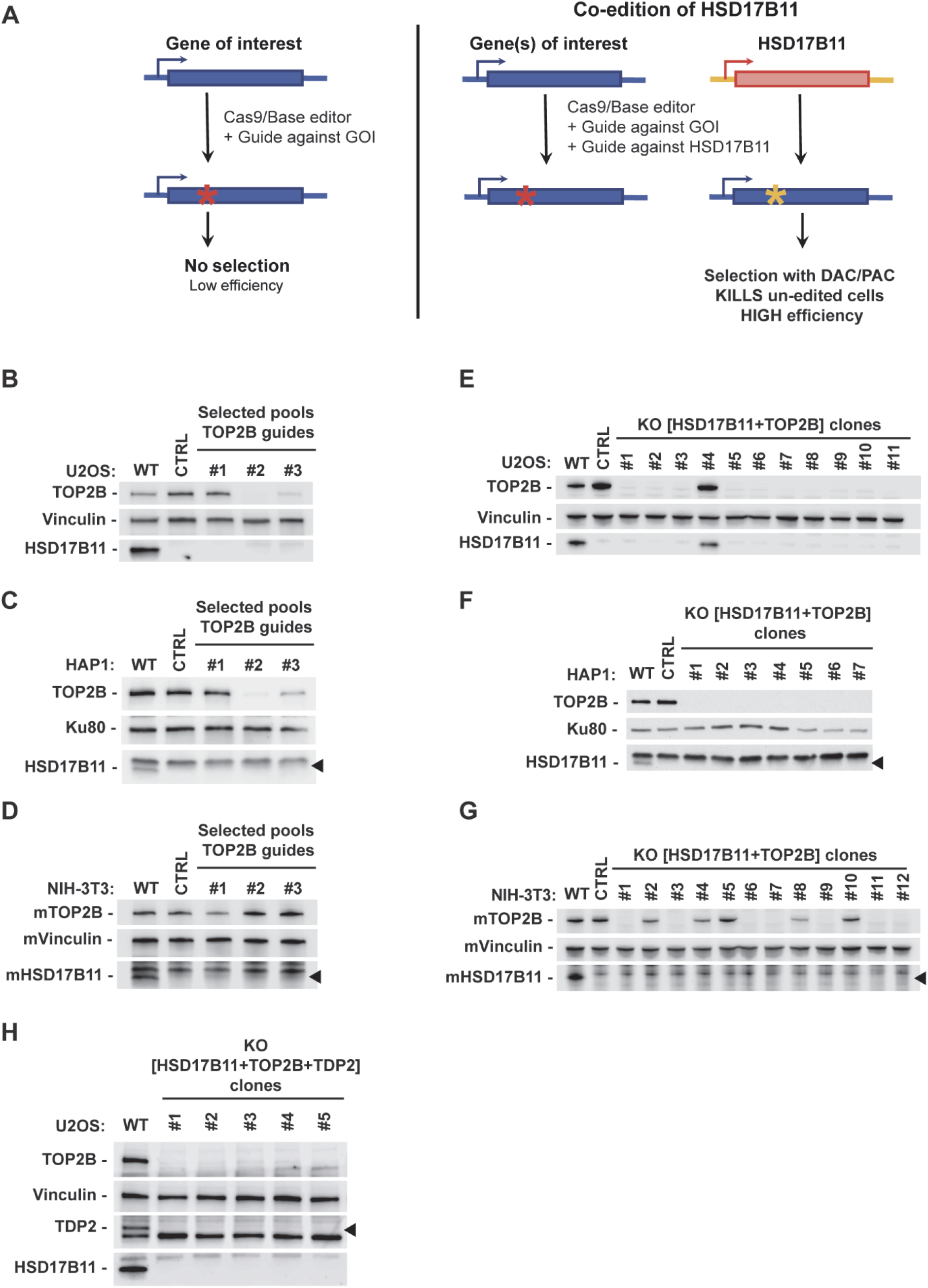
**A**. Scheme depicting the co-inactivation approach used to enrich for successfully edited cells. **B-G**. Analysis by immunoblotting of HSD17B11 and TOP2B levels in U2OS, HAP1 and NIH-3T3 cells that are either WT, CTRL (HSD17B11 knock-out) or co-edited for TOP2B. B, C, and D represent blots of selected populations, used to identify efficient gRNA(s), while E, F and G represent blots of individual clones. To isolate TOP2B KO clones, the gRNAs #2 and #1 were chosen for human and mouse, respectively. **H**. Analysis by immunoblotting of HSD17B11, TOP2B and TDP2 levels in U2OS cells that are either WT or simultaneously co-edited for TOP2B and TDP2. Vinculin or Ku80 was used as a loading control. The black arrows indicate the band corresponding to HSD17B11 or TDP2, as indicated.

Two human cell lines (U2OS and HAP1) and one mouse cell line (NIH-3T3) were transfected with a 1:3 mixture of plasmids co-expressing Cas9 and gRNAs targeting HSD17B11 or the GOI, respectively. Seven days after transfection, cells were selected with 2 µM PAC for human cells or 8 µM PAC for mouse cells. PAC-resistant populations were generated using three different gRNAs targeting TOP2B, and editing efficiency was assessed by immunoblotting (**Fig. 2B-D**). This allowed the identification of gRNA #2 as the most efficient one against human TOP2B in both U2OS and HAP1 cells (**Fig. 2B,C**). The marked reduction in TOP2B levels obtained with this gRNA suggests that the resulting selected populations could potentially be used directly for downstream experiments. In mouse cells, gRNA #1 was the most efficient, although it produced only a partial reduction in TOP2B protein levels (**Fig. 2D**). These results show that PAC selection can be used to rapidly identify efficient gRNAs.

### Co-inactivation of HSD17B11 along with selection with PAC can be exploited to isolate clones lacking detectable expression of one or multiple GOIs

Cell populations generated with the most efficient gRNAs were seeded by limiting dilution to isolate individual clones, which were subsequently screened by immunoblotting (**Fig. 2E-G**). As expected, clones lacking detectable TOP2B expression were identified with high yield in U2OS (10/11, 91%, **Fig. 2E**) and HAP1 (7/7, 100%, **Fig. 2F**) and with intermediate yield in NIH-3T3 (7/12, 59%, **Fig. 2G**).

After validating this approach for the inactivation of a single GOI, we next challenged its efficiency by simultaneously targeting multiple genes. U2OS cells were therefore transfected with a mixture of plasmids co-expressing Cas9 and gRNAs targeting HSD17B11, TOP2B, and TDP2 (ratio 1:1.5:1.5), the latter encoding tyrosyl-DNA phosphodiesterase 2, an enzyme involved in the repair of topoisomerase II-associated DNA damage. Remarkably, all five HSD17B11-inactivated clones also lacked detectable TOP2B and TDP2 protein expression (**Fig. 2H**), demonstrating that PAC selection can also facilitate the isolation of cells carrying simultaneous inactivation of multiple genes.

Altogether, these data show how the HSD17B11/PAC selection can be used to readily i) identify the most active gRNA(s), and ii) isolate clones successfully inactivated for one or multiple GOIs.

### Co-inactivation of HSD17B11 along with selection with PAC allows rapid isolation of successfully edited cells

Having validated the HSD17B11 co-inactivation strategy, we next sought to define the minimal post-transfection time required for the PAC selection to yield the maximum number of clones. Kinetics analysis revealed that selection could be initiated as early as 3 days post-transfection (**Fig. 3**), a time at which resistant colonies begin to form. Notably, by day 4 post-transfection, the number of colonies reached a plateau, corresponding to the maximal yield achievable under the given transfection efficiency. Overall, these results demonstrate that the HSD17B11 co-edition approach enables selection of edited cells as early as 3-4 days after transfection. Considering that PAC treatment only requires 16-24 h to kill HSD17B11 proficient cells, this approach allows enrichment of cells edited for a GOI that is important for cell viability and whose loss would otherwise be diluted in the population with longer post-transfection times.

**Figure 3.**
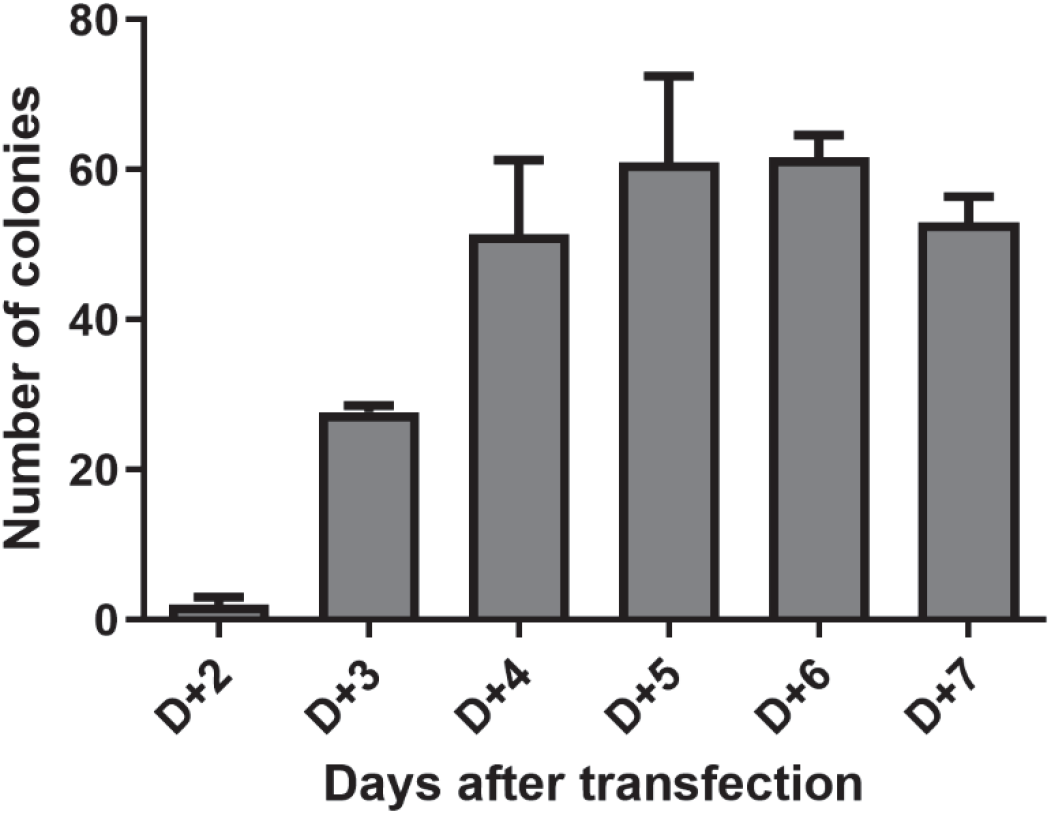
Kinetics of appearance of PAC-resistant clones after transfection. The number of colonies forming at different times post-transfection was evaluated in three independent experiments. Error bars represent standard deviations.

## Discussion

This study provides a framework for exploiting the HSD17B11-dependent bioactivation of PAC into cytotoxic species to rapidly identify efficient CRISPR/Cas9 gRNAs and isolate cells inactivated for a GOI. We anticipate that the same strategy is compatible with the use of Cas12a/Cpf1 or base- and prime editors **(Pacesa et al., 2024)**. Analyzing the resistant cell populations for GOI inactivation provides a rapid assessment of gRNA efficiency, which remains empirical despite continuous progress with the different prediction algorithms. As shown with the human TOP2B gRNA #2, which achieves near-complete depletion of TOP2B protein, cell populations can be directly used for experiments to avoid clonal effects. For less efficient gRNAs and/or for cell lines more difficult to engineer, isolating clones allows recovery of successfully edited cells, as exemplified with the mouse TOP2B gRNA #1 showing low efficiency.

Importantly, we establish here that selection with PAC can be performed as early as 3 days post-transfection, and this duration can probably be shortened by directly transfecting with CRISPR/Cas9 ribonucleoprotein complexes **(Zuris et al., 2015)**. Reducing the post-delivery time can be critical to avoid the progressive depletion of the population from cells inactivated in GOI that is important for cell proliferation/fitness. Mechanistically, DAC and their PAC derivatives are oxidized in a NAD^+^-dependent manner by HSD17B11 into reactive DACones and PACones, which quickly form covalent adducts with multiple intracellular proteins, including essential proteins such as PSMD2, a 19S proteasome subunit **(Demange et al., 2022)**. This results in an acute ER stress, inhibition of the ubiquitin-proteasome system and cell death by apoptosis. This process is rapid, with complete cell death being achieved in less than 16 h **(Bossuat et al., 2023)**. Being able to quickly eliminate unedited cells represents one important benefit of using PAC for selection.

In contrast to ATP1A1, the target of ouabain-based selection, HSD17B11 is non-essential in human cells, and HSD17B11-knockout mice are viable and display only relatively mild phenotypes evocative of increased androgen levels, such as increased heart weight and hyperactivity **(Mouse Genome Informatics, 2024)**. Although HSD17B11 inactivation may affect androgen metabolism *in vivo*, such effects may have limited consequences in cultured cells maintained in complete medium. This is supported by our data showing that HSD17B11 inactivation had no significant effect on doubling time across multiple cell lines. Importantly, ATP1A1 regulates intracellular Na^+^ and K^+^ homeostasis and indirectly influences Ca^2+^ levels, thereby affecting numerous cellular processes, including cell death. Consequently, ATP1A1 mutations, even without impacting on cell viability, could alter cellular responses, a potential limitation that does not apply to the approach described here.

Another advantage of PAC-based selection is that its mechanism of action does not rely on DNA damage induction to achieve cell death, in contrast to other selections, such as YM155 used to select for SLC35F2-inactivated cells **(Kim et al., 2020)**. YM155 is a DNA-damaging agent whose intracellular accumulation relies on the transporter SLC35F2 **(Winter et al., 2014)**. The use of DNA-damaging agents for selection may therefore be undesirable, as it could introduce additional genomic alterations into the cells under study.

It is noteworthy that the HSD17B11-dependent cytotoxicity of DACs and derivatives was previously established in several human cell lines (HAP1, U2OS, A549). The present study extends these observations to three additional widely used human cell lines (HCT116, HEK293T and HeLa), as well as to a murine cell line. We anticipate that this selection principle could be extended to other cells or organisms expressing a functional HSD17B11 orthologue capable of bioactivating PACs. The applicability of PAC selection to mouse cells represents another important advantage over ouabain-based selection, which is poorly suited to rodents because of their intrinsic resistance **(Lingrel, 2010)**.

While HSD17B11 is ubiquitously expressed in human tissues, a few cell lines do not express it. This is the case for several human breast carcinoma cell lines **(Jin et al., 2023; The Human Protein Atlas)**, including T47D **(Demange et al., 2022)**. In this cellular background, prodrugs activated by different enzymes could be exploited to achieve the same goal. For example, we previously identified prodrugs that can be bioactivated by either HSD17B11 or RDH11 and could therefore potentially be exploited for selection **(Demange et al., 2022)**. Like HSD17B11, RDH11 is a non-essential enzyme in human cells **(Arafeh et al., 2025; DepMap, 2026b)** and RDH11 knock-out mice are viable while showing only a delay in visual adaptation to the dark, in agreement with a role in ocular retinol metabolism **(Kasus-Jacobi et al., 2005)**. Another attractive enzyme would be the catechol-O-methyl transferase (COMT) that bioactivates EAPB02303 into EAPB04303, a potent inhibitor of microtubule polymerization **(Bigot et al., 2025)**. However, COMT inactivation seems to reduce the fitness of several cell lines **(DepMap, 2026a)**. Such alternative enzyme/prodrug pairs could also enable sequential selection for the inactivation of additional genes in cells in which HSD17B11 has already been disrupted. A limitation of such sequential strategies is the need for appropriately matched control cells carrying the same inactivation of all genes used for selection.

Finally, an important practical advantage demonstrated in this study is the possibility of using readily accessible racemic PACs rather than enantiopure compounds, which are more difficult and costly to produce. This takes advantage of the relative innocuity of the inactive enantiomer which is not oxidized by HSD17B11. It is important to note that, while this study uses the PAC depicted in **Fig. 1A** (1-(4-octylphenyl)penta-1,4-diyn-3-ol)), a wide variety of synthetic DAC analogues could be exploited similarly.

## Acknowledgements

This work was funded by the Région Occitanie “Prostacure” pre-maturation program, the CNRS pre-maturation program “Easy-Crisp”, the french National Research Agency (ANR-24-CE44-1817-03 “TOP2-G4”), and by the TIRIS-Scaling-up science “SDR-to-lead” program supported by the Région Occitanie, the European Development Fund (ERDF), and the French government, through the France 2030 project managed by the National Research Agency (ANR) with the reference number “ANR-22-EXES-0015”.

## Author contributions

S.Br. and Y.G. conceived the study and acquired funding. S.Br., Y.G., S.Ba. and B.D. supervised the students. S.Br., B.D. and M.B. designed the experiments. S.Br. and B.D. wrote the original draft. S.Br., B.D. and M.B. prepared the figures and analyzed the data. B.D., M.B., A.R., S.G., L.F., M.C., P.F., D.G. and P.S. performed the experiments. P.S., S.Ba., V.M., V.B.- G. and Y.G. devised an improved synthetic route to PAC. P.S. and S.Ba. produced PAC. All authors reviewed and approved the final manuscript.

## Data availability statement

All data generated or analyzed during this study are included in this published article.

## Competing Interests Statement

There are no conflicts of interest to report.

## Methods

### Cell lines

U2OS (ATCC), HCT-116 (Horizon Discovery), HEK293T (ATCC), HeLa clone I3 (gift from Titia de Lange) and NIH-3T3 (gift from Florence Larminat) cells were grown in DMEM 10% FBS; HAP-1 (Horizon Discovery) in IMDM 10% FBS. All cell media contained penicillin and streptomycin (Thermo Fisher Scientific) and cells were grown at 37 °C in a 5% CO_2_ humidified incubator. Cells were used at low passage and routinely confirmed free of Mycoplasma. Cells were treated in complete growth medium.

### Chemicals

The PAC, synthesized as previously described **(Bossuat et al., 2025; Bossuat et al., 2023)**, was resuspended at 5 mM in DMSO and added directly to complete medium for selections.

### Plasmid transfection and stable cell line generation

Cells were transfected at 90% confluency in 60 mm dishes using 5 µg DNA and lipofectamine 2000 (Thermo Fisher Scientific) following manufacturer’s instructions. Cells were transfected with pCAG-eSpCas9-2A-GFP plasmid (Addgene Plasmid #79145) coding for the *S. pyogenes* Cas9 K848A K1003A R1060A variant and co-expressing a previously validated gRNA against human HSD17B11 (target TGTAATCAGCACGATTTCGC, **(Demange et al., 2022)**, Addgene Plasmid # 161923), or against mouse HSD17B11 (target sequence GGTGATCAGGACGATCTCTC) for NIH3T3 cells. For co-edition of TOP2B, co-transfection was performed with the same plasmid but expressing a gRNA against human TOP2B (target sequences: #1 TGATACATATATTGGGTCAG, #2 GTCAGTGGAGCCATTGACGC, #3 GCAATAATTACCTGCGTCAA), human TDP2 (target sequence GGCATCGCAGCTTGCGACCG) or mouse TOP2B (target sequences: #1 AGATGTAGGGATGAACTGCA, #2 TGATACATACATTGGATCAG, #3 CAATGTATGTATCAGGACGA). gRNAs were designed using Benchling. For isolating clones, gRNAs #2 and #1 were chosen respectively for human and mouse TOP2B. Six days after transfection, 250 000 cells were seeded in large dishes and selected the day after with a toxic concentration of PAC (2 and 8 µM for human and mouse cells, respectively). Individual clones were isolated, and gene inactivation was validated by immunoblotting.

### Immunoblotting

For whole-cell extracts (WCE), cells were washed with cold PBS and scraped in 75 µL SDS-lysis buffer (120 mM Tris-HCl pH 6.8, 20% glycerol, 4% SDS), incubated 5 min at 95 °C and passed 10 times through a 25 G needle. Protein concentration was measured by absorbance at 280 nm using NanoDrop spectrometer (Thermo Fisher Scientific) and, after adjustment with SDS-Lysis buffer, extracts were diluted by addition of an equal volume of SDS-Loading Buffer (5 mM Tris pH 6.8, 0.01% bromophenol blue, 0.2 M dithiothreitol). Immunoblotting was performed with 25-50 µg of WCE per condition, along with the PageRuler Prestained Protein Ladder or PageRuler Plus Prestained Protein Ladder (Invitrogen). Proteins were separated on gradient gels (BioRad 4-12% TGX or NuPAGE Bis-Tris 4-12% midi pre-cast gels) and transferred onto Protran 0.45 µm nitrocellulose membranes (GE Healthcare). Homogeneous loading and transfer were checked by Ponceau S staining. When necessary, membranes were cut into horizontal strips to simultaneously probe for multiple proteins. Membranes were blocked with PBS, 0.1% Tween-20 (Sigma-Aldrich) (PBS-T buffer) containing 5% non-fat dry cow milk, washed and incubated overnight at 4 °C with primary antibodies. After extensive washes, membranes were probed 1 h at room temperature with HRP-conjugated goat secondary antibodies (Jackson Immunoresearch Laboratories) diluted 1/10000 in PBS-T. After incubation with peroxidase chemiluminescent substrates (BioRad Clarity or Advansta WesternBright ECL), membranes were exposed using autoradiographic films or scanned using an imager (ChemiDoc Touch, Biorad or Odyssey XF, LI-Cor Biosciences). The following primary antibodies were used: mouse monoclonal antibodies anti-Vinculin (clone 7F9, Santa Cruz Biotechnologies, SC-73614, 1/1000) and anti-Ku80 (clone 111 ; Thermo Fisher Scientific) ; rabbit polyclonal antibodies anti-TOP2B (Bethyl Laboratories, A300-950A-1, 1/1000), anti-HSD17B11 (Invitrogen, PA5-54469, lot A117963 and 000053616, 1/500 for **Fig. 1B,2B,2E,2D,2G** or Proteintech, 16303-1-AP, lot 00007618, 1/250, for **Fig. 2C,F**, revealing two bands in human cells, the lower being HSD17B11), polyclonal antibody anti-TDP2 (Bethyl Laboratories, A302-737A, 1/1000, for **Fig.2H**, revealing two bands in human cells, with the upper one being TDP2). All primary antibodies were diluted in PBS-T containing 1% bovine serum albumin (immunoglobulin- and lipid-free fraction V BSA, Sigma-Aldrich) and 0.02% sodium azide as a preservative.

### Viability assays

Cell viability was analyzed using SRB assays. Cells were seeded in 96-well plates 24 h before being treated continuously for 72 h with the indicated concentration of PAC. For analysis, cells were fixed for 1 h at 4 °C by addition of cold trichloroacetic acid at a 3.33% final concentration. After being washed four times with water and dried, cells were stained by a 30 min incubation in a solution of 0.057% (wt:vol) SRB in 1% acetic acid. The wells were washed four times with 1% acetic acid, dried and the dye was resuspended by a 1-2 h incubation in a 10 mM Tris-Base solution. Absorbance at 490 nm of each well was measured (µQuant plate reader, Bio-tek) and used as a readout of cell number. For calculation, background absorbance was subtracted to each value and the data were normalized to the value measured in untreated wells. Each point was measured in duplicate and the graphs correspond to at least three independent experiments. IC_50_ were computed with the GraphPad Prism software using a non-linear regression to a curve fit (log[inhibitor] *vs* normalized response; variable slope). The error bars represent standard deviation (SD). The resistance factor was defined as the ratio of the IC_50_ value obtained in HSD17B11 knock-out cells to that measured in wild-type (WT) cells. The concentration of selection was determined as the concentration required to kill all unmodified cells while preserving most cells lacking the HSD17B11 gene.

### Cell proliferation assays

Cells were seeded at densities optimized for each cell line in three distinct 60 mm dishes: 100 000 cells for HEK293T, U2OS, HeLa, NIH3T3; 200 000 cells for HAP1, HCT116. Each day, one dish was processed. Cells were washed with PBS and collected by trypsination and counted twice. This procedure was repeated daily for up to 4 days post-seeding. Because U2OS cells reached confluency, their proliferation was calculated over a 3-day period. Doubling time of each cell line was computed with GraphPad Prism software using a non-linear regression (exponential growth equation). The experiment was conducted three times independently.

### PAC clonogenic assays after transfection

Cells were transfected at 90% confluency in 60 mm dishes using Lipofectamine 2000 and 5 µg of pCAG-eSpCas9-2A-GFP plasmid co-expressing a gRNA targeting human *HSD17B11*(Addgene plasmid #161923). Twenty-four hours post-transfection, cells were maintained in a seeding scheme to enable selection treatment from day 2 to day 7 post-transfection. Cells were either seeded at 5 × 10^5^ cells in 140 mm dish for treatment the following day or maintained at appropriate densities to allow subsequent reseeding steps. Each day, one 140 mm dish was treated with 2 µM of PAC, generating a continuous treatment time course across the indicated period. After 2-3 days selection, the medium was replaced and cells were maintained in culture until PAC-resistant clones appeared. Once colonies reached sufficient size, cells were fixed with 3.33% cold trichloroacetic acid for 1 h at 4 °C. After being washed four times with water and air dried, cells were stained by a 30 min incubation in a solution of 0.057% (wt:vol) SRB in 1% acetic acid at RT. The dishes were washed four times with 1% acetic acid and air dried. Colonies were counted in each 140 mm dish to assess the kinetics of PAC-resistant clone appearance. The experiment was conducted three times independently, and data were analyzed using GraphPad Prism software. The error bars represent standard deviation (SD).

## Notes

### Competing Interest Statement

The authors have declared no competing interest.

### Summary of Updates

Additional experiments have been performed (Figure 2H) by a Luce Fontanie--Roger (new author), the abstract and text have been corrected (grammar, spelling and missing information), and additional funding sources have been included.

## References

Agudelo, D., Duringer, A., Bozoyan, L., Huard, C.C., Carter, S., Loehr, J., Synodinou, D., Drouin, M., Salsman, J., Dellaire, G., et al. (2017). Marker-free coselection for CRISPR-driven genome editing in human cells. Nat Methods 14, 615–620, doi:10.1038/nmeth.4265.

Arafeh, R., Shibue, T., Dempster, J.M., Hahn, W.C., and Vazquez, F. (2025). The present and future of the Cancer Dependency Map. Nat Rev Cancer 25, 59–73, doi:10.1038/s41568-024-00763-x.

Bigot, K., Patinote, C., Garambois, V., Chouchou, A., Gayraud-Paniagua, S., Vie, N., Maggipinto, Y., Smyej, E., Robin, M., Machu, M., et al. (2025). Inhibiting microtubule polymerization with EAPB02303, a prodrug activated by catechol-O-methyl transferase, enhances paclitaxel effect in pancreatic cancer models. Cell death & disease 16, 441, doi:10.1038/s41419-025-07747-1.

Billon, P., Bryant, E.E., Joseph, S.A., Nambiar, T.S., Hayward, S.B., Rothstein, R., and Ciccia, A. (2017). CRISPR-Mediated Base Editing Enables Efficient Disruption of Eukaryotic Genes through Induction of STOP Codons. Mol Cell 67, 1068–1079 e1064, doi:10.1016/j.molcel.2017.08.008.

Biondo, E.D., Spontarelli, K., Ababioh, G., Mendez, L., and Artigas, P. (2021). Diseases caused by mutations in the Na(+)/K(+) pump alpha1 gene ATP1A1. Am J Physiol Cell Physiol 321, C394–C408, doi:10.1152/ajpcell.00059.2021.

Bossuat, M., Preuilh, N., Seigneur, P., Fabing, I., Pradel, C., Peixoto, A., Maraval, V., Bernardes-Genisson, V., Ballereau, S., Britton, S., et al. (2025). A plug and play approach to structural variations of bio-inspired anticancer lipidic phenyl dialkynylcarbinols. RSC Adv 15, 38307–38320, doi:10.1039/d5ra06473b.

Bossuat, M., Rulliere, P., Preuilh, N., Peixoto, A., Joly, E., Gomez, J.G., Bourkhis, M., Rodriguez, F., Goncalves, F., Fabing, I., et al. (2023). Phenyl dialkynylcarbinols, a Bioinspired Series of Synthetic Antitumor Acetylenic Lipids. J Med Chem 66, 13918–13945, doi:10.1021/acs.jmedchem.3c00859.

Brereton, P., Suzuki, T., Sasano, H., Li, K., Duarte, C., Obeyesekere, V., Haeseleer, F., Palczewski, K., Smith, I., Komesaroff, P., et al. (2001). Pan1b (17betaHSD11)-enzymatic activity and distribution in the lung. Mol Cell Endocrinol 171, 111–117, doi:10.1016/s0303-7207(00)00417-2.

Contreras, R.G., Torres-Carrillo, A., Flores-Maldonado, C., Shoshani, L., and Ponce, A. (2024). Na(+)/K(+)-ATPase: More than an Electrogenic Pump. Int J Mol Sci 25, doi:10.3390/ijms25116122.

Croyle, M.L., Woo, A.L., and Lingrel, J.B. (1997). Extensive random mutagenesis analysis of the Na+/K+-ATPase alpha subunit identifies known and previously unidentified amino acid residues that alter ouabain sensitivity--implications for ouabain binding. Eur J Biochem 248, 488–495, doi:10.1111/j.1432-1033.1997.00488.x.

Demange, P., Joly, E., Marcoux, J., Zanon, P.R.A., Listunov, D., Rulliere, P., Barthes, C., Noirot, C., Izquierdo, J.B., Rozie, A., et al. (2022). SDR enzymes oxidize specific lipidic alkynylcarbinols into cytotoxic protein-reactive species. eLife 11, doi:10.7554/eLife.73913.

DepMap, B.I. (2026a). COMT DepMap Gene Summary (Broad Institute).

DepMap, B.I. (2026b). RDH11 DepMap Gene Summary (Broad Institute).

DepMap, B.I. (2026c). TOP2B DepMap Gene Summary (Broad Institute).

Djamshidi, M., Hill, A., Heshmatzad, K., Langley, J., Krowicki, H., Ali, M., Yang, Y., Tanida, R., Abdul-Careem, M.F., Billon, P., et al. (2026). FAME-CRISPR improves CRISPR-Cas9 genome editing via HDAC inhibition and engineered virus-like particle delivery. Cell Rep Methods 6, 101248, doi:10.1016/j.crmeth.2025.101248.

Gagnon, K.B., and Delpire, E. (2020). Sodium Transporters in Human Health and Disease. Front Physiol 11, 588664, doi:10.3389/fphys.2020.588664.

Horiguchi, Y., Araki, M., and Motojima, K. (2008). Identification and characterization of the ER/lipid droplet-targeting sequence in 17beta-hydroxysteroid dehydrogenase type 11. Arch Biochem Biophys 479, 121–130, doi:10.1016/j.abb.2008.08.020.

Jin, H., Zhang, C., Zwahlen, M., von Feilitzen, K., Karlsson, M., Shi, M., Yuan, M., Song, X., Li, X., Yang, H., et al. (2023). Systematic transcriptional analysis of human cell lines for gene expression landscape and tumor representation. Nat Commun 14, 5417, doi:10.1038/s41467-023-41132-w.

Kasus-Jacobi, A., Ou, J., Birch, D.G., Locke, K.G., Shelton, J.M., Richardson, J.A., Murphy, A.J., Valenzuela, D.M., Yancopoulos, G.D., and Edwards, A.O. (2005). Functional characterization of mouse RDH11 as a retinol dehydrogenase involved in dark adaptation in vivo. J Biol Chem 280, 20413–20420, doi:10.1074/jbc.M413789200.

Kim, K.T., Park, J.C., Jang, H.K., Lee, H., Park, S., Kim, J., Kwon, O.S., Go, Y.H., Jin, Y., Kim, W., et al. (2020). Safe scarless cassette-free selection of genome-edited human pluripotent stem cells using temporary drug resistance. Biomaterials 262, 120295, doi:10.1016/j.biomaterials.2020.120295.

Levesque, S., Mayorga, D., Fiset, J.P., Goupil, C., Duringer, A., Loiselle, A., Bouchard, E., Agudelo, D., and Doyon, Y. (2022). Marker-free co-selection for successive rounds of prime editing in human cells. Nat Commun 13, 5909, doi:10.1038/s41467-022-33669-z.

Lingrel, J.B. (2010). The physiological significance of the cardiotonic steroid/ouabain-binding site of the Na,K-ATPase. Annu Rev Physiol 72, 395–412, doi:10.1146/annurev-physiol-021909-135725.

Lyu, P., Javidi-Parsijani, P., Atala, A., and Lu, B. (2019). Delivering Cas9/sgRNA ribonucleoprotein (RNP) by lentiviral capsid-based bionanoparticles for efficient ‘hit-and-run’ genome editing. Nucleic Acids Res 47, e99, doi:10.1093/nar/gkz605.

Mouse Genome Informatics (2024). Hsd17b11<tm1.1(KOMP)Vlcg> Targeted Allele Detail (The Jackson Laboratory).

Nambiar, T.S., Baudrier, L., Billon, P., and Ciccia, A. (2022). CRISPR-based genome editing through the lens of DNA repair. Mol Cell 82, 348–388, doi:10.1016/j.molcel.2021.12.026.

Pacesa, M., Pelea, O., and Jinek, M. (2024). Past, present, and future of CRISPR genome editing technologies. Cell 187, 1076–1100, doi:10.1016/j.cell.2024.01.042.

Ran, F.A., Hsu, P.D., Wright, J., Agarwala, V., Scott, D.A., and Zhang, F. (2013). Genome engineering using the CRISPR-Cas9 system. Nat Protoc 8, 2281–2308, doi:10.1038/nprot.2013.143.

Slaymaker, I.M., Gao, L., Zetsche, B., Scott, D.A., Yan, W.X., and Zhang, F. (2016). Rationally engineered Cas9 nucleases with improved specificity. Science 351, 84–88, doi:10.1126/science.aad5227.

The Human Protein Atlas. Cell line - HSD17B11 (The Human Protein Atlas).

Uuskula-Reimand, L., and Wilson, M.D. (2022). Untangling the roles of TOP2A and TOP2B in transcription and cancer. Sci Adv 8, eadd4920, doi:10.1126/sciadv.add4920.

Winter, G.E., Radic, B., Mayor-Ruiz, C., Blomen, V.A., Trefzer, C., Kandasamy, R.K., Huber, K.V.M., Gridling, M., Chen, D., Klampfl, T., et al. (2014). The solute carrier SLC35F2 enables YM155-mediated DNA damage toxicity. Nat Chem Biol 10, 768–773, doi:10.1038/nchembio.1590.

Zuris, J.A., Thompson, D.B., Shu, Y., Guilinger, J.P., Bessen, J.L., Hu, J.H., Maeder, M.L., Joung, J.K., Chen, Z.Y., and Liu, D.R. (2015). Cationic lipid-mediated delivery of proteins enables efficient protein-based genome editing in vitro and in vivo. Nat Biotechnol 33, 73–80, doi:10.1038/nbt.3081.

