## Supplementary Figure 1 for "Exploiting HSD17B11-dependent dialkynylcarbinols cytotoxicity for facile CRISPR/Cas9-based gene inactivation"

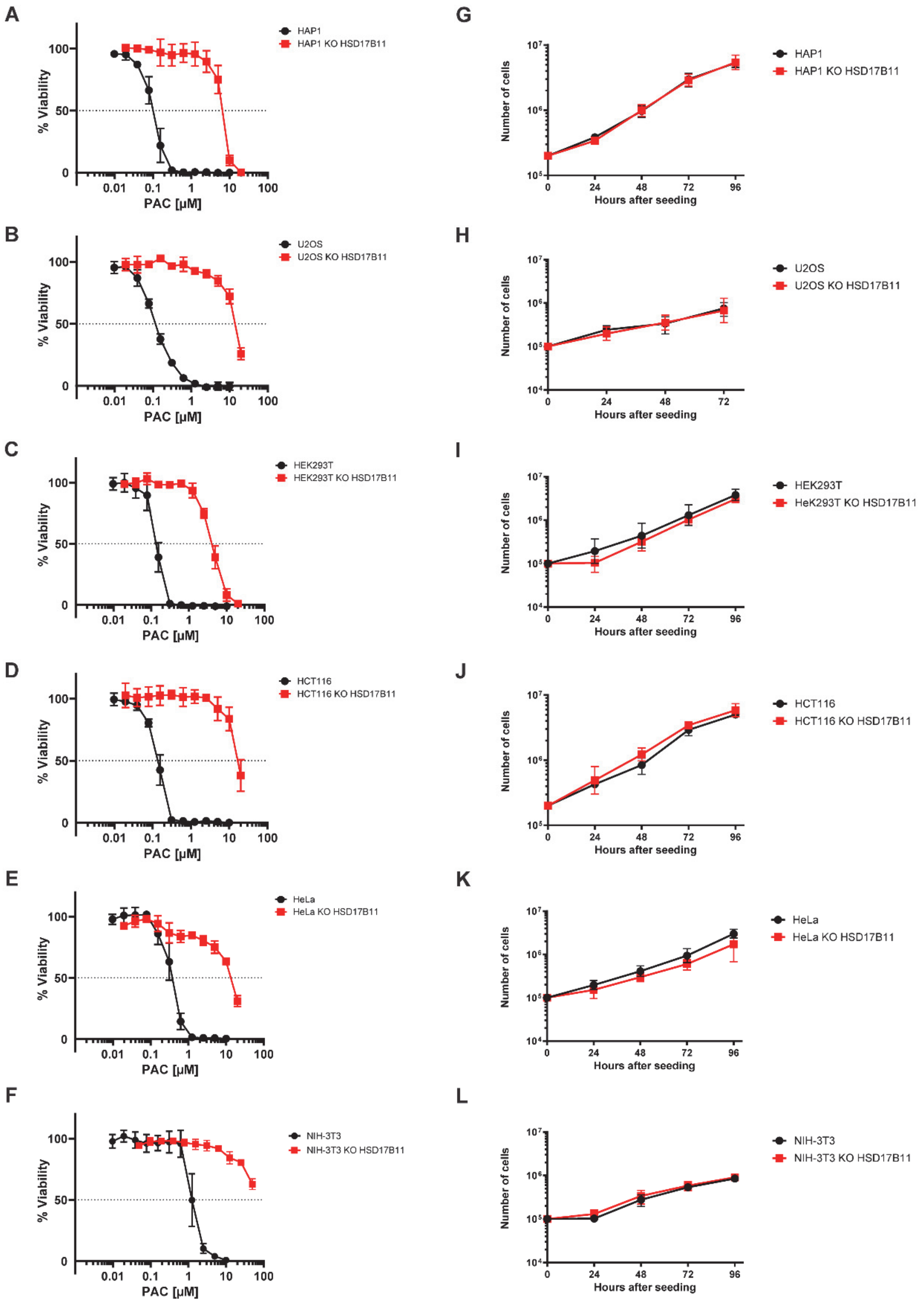


**Figure S1: A-F** Cell viability analysis of each cell line, including WT and an isolated clone for HSD17B11 knock-out treated for 72 h with the indicated concentrations of PAC. **G-L** Proliferation curves of each cell line, including WT and an isolated HSD17B11 knock-out clone.
